# Diatom phytochromes integrate the entire visible light spectra for photosensing in marine environments

**DOI:** 10.1101/2023.01.25.525482

**Authors:** Carole Duchêne, Jean-Pierre Bouly, Juan José Pierella Karlusich, Julien Sellés, Benjamin Bailleul, Chris Bowler, Maurizio Ribera d’Alcalà, Angela Falciatore, Marianne Jaubert

**Affiliations:** CNRS, Sorbonne Université, Institut de Biologie Physico-Chimique, Laboratoire de Biologie du chloroplaste et perception de la lumière chez les microalgues, UMR7141, F-75005 Paris, France; Institut de Biologie de l’ENS (IBENS), Département de Biologie, École Normale Supérieure, CNRS, INSERM, Université PSL, 75005 Paris, France; Stazione Zoologica Anton Dohrn, 80121 Naples, Italy

## Abstract

Aquatic life is strongly structured by light gradients, with gradual decrease in light intensity and differential attenuation of sunlight wavelengths with depth. How phytoplankton perceive these variations is unknown. By providing the first *in vivo* quantitative assessment of the action of marine diatom phytochrome photoreceptors (DPH), we show that they efficiently trigger photoreversible responses across the entire light spectrum, unlike current models of phytochrome photosensing. The distribution and activity of DPHs in the environment indicate that they are extremely sensitive detectors of spectral light variations related to depth and optical properties of the water column in temperate and polar oceans, revealing a completely novel view of how light is perceived in the marine environment.

## Main Text

Continuous perception of the environment is an essential need for all organisms. Signals associated with changes in light (intensity, color, duration and direction) are among the most informative. Accordingly, photoreceptors have evolved with various degrees of complexity to perceive and discriminate information associated with light signals. Among them, phytochromes are red/far-red light-sensing proteins with multiple roles in land plants, also present in photosynthetic and non-photosynthetic bacteria, fungi and diverse algae *(1)*. These photoreceptors are dimeric chromoproteins, which covalently bind to a tetrapyrrole chromophore. They typically convert, photoreversibly, between a red (R)-absorbing form (Pr) and a far-red (FR)-absorbing form (Pfr), and the resulting equilibrium between the two forms under a given light field modulates the triggered biological responses *(2)*. Some algal phytochromes from prasinophytes, glaucophytes and brown algae exhibit orange-yellow/FR, R/blue (B) or FR/green (G) photocycles, respectively *(3, 4)*. As R and FR bands are strongly attenuated by water *(5*), these shifts have been interpreted as an adaptive spectral tuning of algal phytochromes to aquatic light fields *(3, 4)*. However, this is clearly not a general feature because phytochromes from marine diatom microalgae (DPH) do display R/FR photocycles *(6)*.Moreover, in the model diatom *Phaeodactylum tricornutum*, exposure to FR light leads to gene expression changes which are lost in DPH knock-out mutants (*dph*), suggesting that the Pr form is biologically active. This led to the hypothesis that DPH could be involved in the sensing of R/FR from sunlight at the surface or from indirect sources such as chlorophyll autofluorescence or Raman scattering at depth *(6)*.

To explore the ecological and functional significance of DPH-based light sensing, we first investigated the biogeographical distribution of DPH-containing diatoms by looking at the presence of *DPH* genes in available genomes and transcriptomes of cultivated species and in *Tara* Oceans environmental sequences *(7, 8)*. While diatoms are found in all sampled regions, DPH-containing diatoms (mostly centric diatoms from the Thalassiosirales and Cymatosirales order in the environmental data) are distributed with a strong latitudinal pattern, with *DPH* genes essentially being excluded from tropical regions (Fig.1A, Fig.S1). Correlation analysis and generalized additive models (Fig.1B and Fig.S1) suggest that these diatoms are present in regions characterized by low temperature and light penetration, and high seasonal variations of chlorophyll. This appears to be a DPH-specific pattern as the diatom B-light photoreceptor aureochromes are ubiquitously distributed and do not show any latitudinal gradient (Fig.S1). Remarkably, recombinant DPH from two genes abundant in the *Tara* Oceans dataset and from five coastal centric diatom species show conserved absorption spectra (Fig.1C and D, Fig. S2, Table S1 and S2), with peaks in FR at 767 nm (+/-2nm, Pfr peak) and in R at 680 nm (+/-4 nm, Pr peak), as previously found for PtDPH *(6)*.

**Fig. 1.**
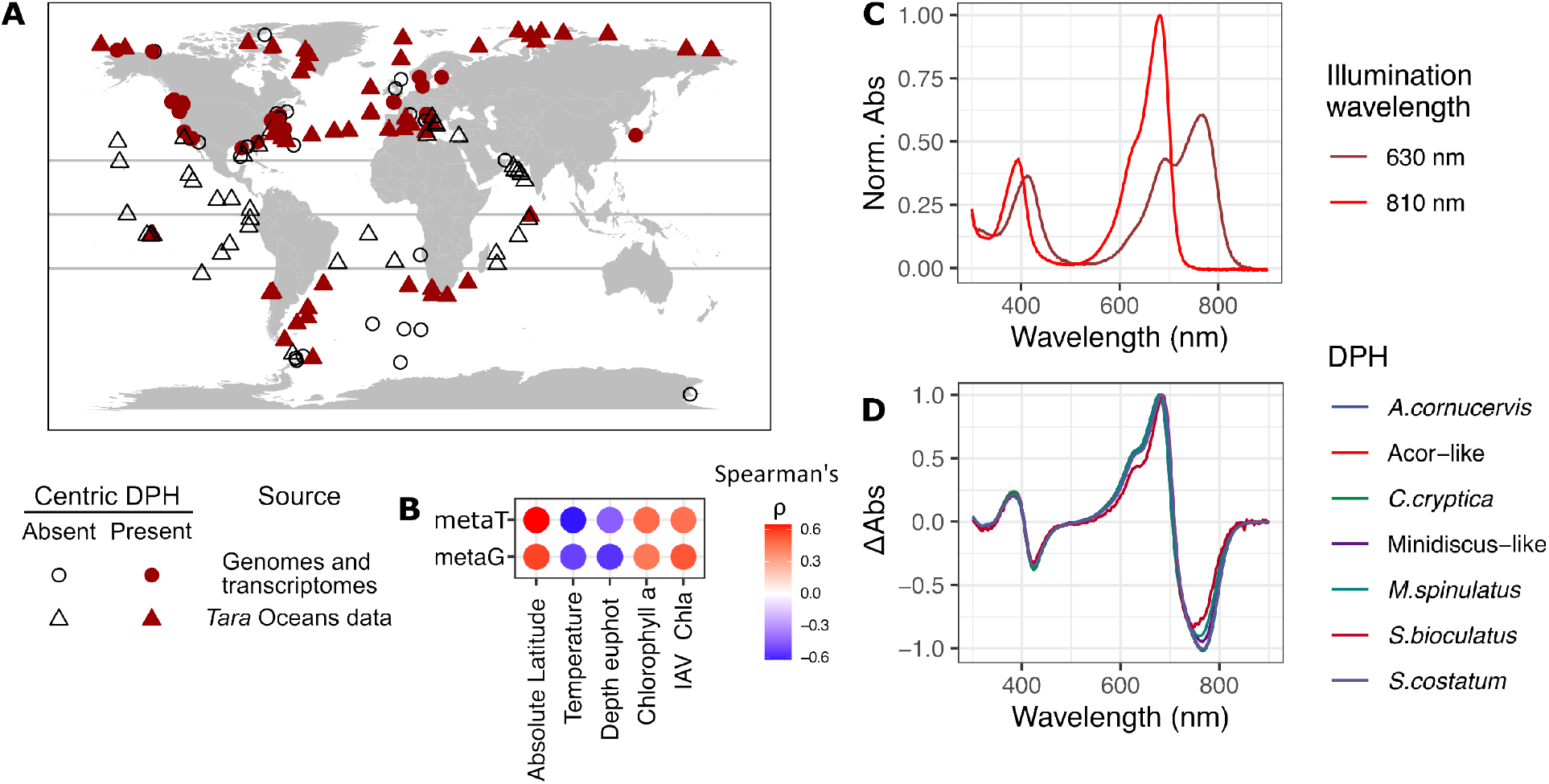
DPHs show a strong latitudinal distribution pattern and conserved spectral properties. (**A**) Map of the presence and absence of centric *DPH* genes based on *Tara* Oceans sequence data and sampling location of centric diatom strains with and without DPH. Equator and Tropics of Cancer and Capricorn are shown. (**B**) Spearman correlation analysis between DPH gene abundance in metagenomic (metaG) read or *DPH* transcript abundance in metatransciptomic (metaT) reads, relative to all centric diatom genes or transcripts, and environmental parameters measured during the *Tara* Oceans campaign (all correlations are significant, p<0.05). Euphot depth: depth of the euphotic zone, i.e. at which the light intensity is 1% of the surface irradiance; IAV Chla: intra annual variation of chlorophyll a concentration measured by satellite. (**C**) Normalized absorption spectra of far-red light illuminated (810nm, red line) and red light illuminated (630nm, darker red line) purified recombinant photosensory module of *Arcocellulus cornucervis–like* (Acor-like) DPH, representing a synthetic environmental gene, expressed in *E. coli* with biliverdin as chromophore. (**D**) Normalized differential absorption spectra of DPH from synthetic environmental genes (“like”) and cloned from cultured diatoms (italic; *Arcocellulus cornucervis Minidiscus spinulatus, Cyclotella cryptica, Shionodiscus bioculatus, Skeletonema costatum*), expressed and illuminated as above.

To solve the apparent incongruence between the DPH absorption spectra and the R-poor light field that diatoms are exposed to, we characterized the wavelength- and intensity-dependence of DPH responses *in vivo*. To this aim, we generated reporter lines in the model species *P. tricornutum* in which the Yellow Fluorescent Protein (YFP) was expressed under control of the DPH-regulated, FR-inducible *HSF4.6a* gene promoter *(6)* in wild type (WT) and *DPH* knock-out genetic backgrounds (Fig. S3A). DPH-dependent fluence rate response curves were obtained from reporter lines exposed for 10 min to different intensities of various monochromatic lights. All wavelengths tested triggered responses, from B to very long FR (Fig 2A), in a DPH-dependent manner, as evidenced using *dph* reporter lines (Fig.S3B). DPH-dependent responses to B, in addition to FR, were also shown by RT-qPCR for *HSF4.6a* and other PtDPH-regulated genes (Fig.S4) *(6)*. We furthermore tested whether DPH responses were photoreversible by exposing the reporter lines first to 10 min of saturating FR light (800 nm) followed by 10 min of different intensities of monochromatic lights (Fig.2A bottom panel). The fluence rate response curves for inhibition showed the same shape in an opposite orientation to the induction experiments, indicating that R (630 and 690 nm) but also G (520 nm), B (405 nm) and FR (730 nm) lights were able to strongly revert DPH responses. In addition, the response induced by saturating bichromatic illuminations show intermediate YFP levels, as evidenced by plotting YFP levels as a function of Pr/Pfr absorbance ratio (Fig. 2B). Taken together, these results indicate that DPH triggers photoreversible responses and perceives waveband ratio not only for R/FR but over the entire visible spectrum, in contrast to what is currently known about phytochromes. Indeed, phytochrome-dependent blue light responses are known in fungi and to some extend in plants *(9–12)*, but photoreversibility over the entire visible light spectra has, to our knowledge, never been shown but it is tempting to speculate it could apply to other phytochromes.

**Fig. 2.**
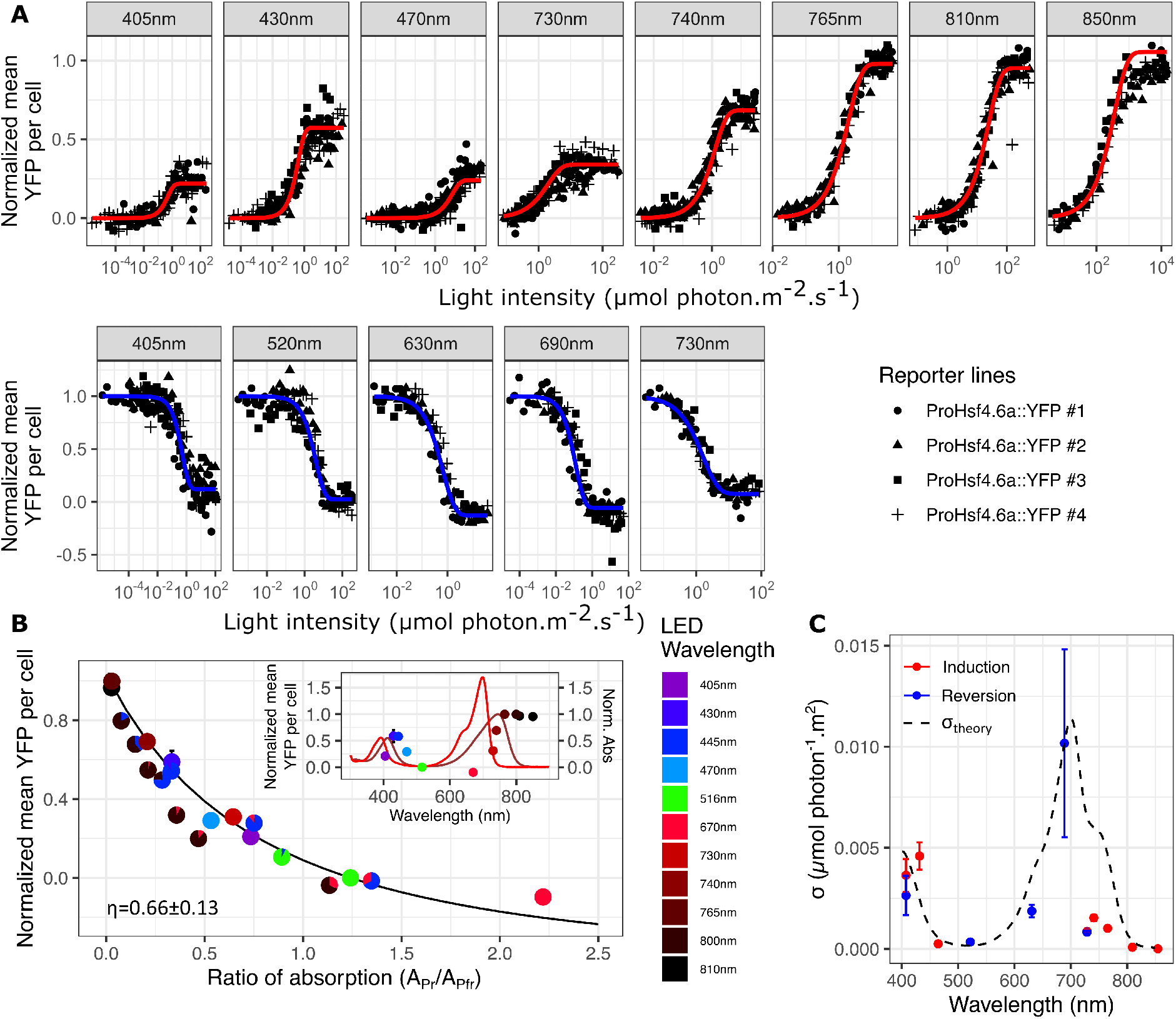
DPH responds to the entire visible light spectra *in vivo*. (**A**) DPH-dependent fluence rate responses (induction, upper panel, and reversion, lower panel) in four ProHsf4.6a::YFP wild type reporter lines. For induction, green-light adapted cells were irradiated for 10 min with light of different colors and intensities. For the reversion response, cells were first exposed to 10 min saturating FR, then to different light treatments for 10 min. Mean YFP per cell were normalized between value in the dark and value after exposure to saturating FR light. (**B**) Plot showing YFP signal at saturating monochromatic lights (from A) and response to mixed wavelengths, used to estimate the ratio of photoconversion yield, η (black curve, Equation 6 in Supplementary Text). Pie charts indicate the relative intensity of each LED color. Inset, YFP signal at saturation superimposed to normalized absorption spectra of PtDPH pure Pfr (dark red) and Pr (red) forms *(6)*. (**C**) Sensitivity of PtDPH to different wavelengths, i.e., the phytochrome photoconversion cross-section σ, resulting from the fitting of Equation 7 in Supplementary Text (colored lines in A). Dashed line represents the theoretical σ, calculated from the absorption spectra of the recombinant protein and η. Values in (B) and (C) are means ±se from the four independent WT lines in (A).

To precisely quantify the effect of different wavelengths and intensities on DPH action, we considered the YFP signal as a direct measure of the percent of DPH in active form (PrPr) and fitted a classical mathematical model for DPH dimer dynamics on the fluence response curves (*(13)* and supplementary text). By fitting the YFP level obtained under saturating monochromatic illuminations as a function of Pr/Pfr absorbance, we estimated the ratio of quantum yields of the Pr→Pfr over Pfr→Pr reactions to be 0.66±0.13 (Fig.2 B). This value is very similar to the one determined from recombinant proteins *in vitro* (0.753, Table S2). We additionally estimated the sensitivity of the DPH response, which should reflect the photoconversion cross-section of both Pr and Pfr (σ, see Supplementary text). Globally, the sensitivity matches with Pr and Pfr absorption spectra with peaks in B, R and FR lights (Fig.2C), but, surprisingly, FR light appears less efficient compared to other wavelengths. This decreased FR sensitivity could be due to different events, e.g., light distortion inside the cell *(14, 15)*, an energetic barrier for low energy FR photons, or signaling events following DPH light absorption in FR compared to other wavebands *(16)*. This is also observed with *dph* reporter lines complemented with the *Thalassiosira pseudonana* DPH, which is more closely related to the DPH found in the open ocean from centric species (Fig.S5, DataS1).

We then explored DPH function *in situ*, based on its novel *in vivo* photoproperties combined with the specificities of the underwater light field. We first projected DPH equilibrium at saturation (% of DPH in PrPr form) in different measured light environments. In contrast to previous projections using R/FR bands only *(6)*, %PrPr increases with depth in a biphasic manner, rapidly in the first 15 m and more gradually below, due to the rapid attenuation of the R band and progressive enrichment in the more penetrative B band (Fig.3A and B). Using both *in situ* and modelled light field data (which we used to explore a wide range of light field scenarios), the presence of algal cells and their associated material, which absorb strongly in the B waveband, was found to reduce %PrPr at depth (Fig.3A, Fig.S6). To a minor extent, DPH equilibrium also increased at low solar elevation, which reflects time of day (dawn and dusk) and latitude (Fig.S6). As evidenced by removing the FR bands from the natural light field, the contribution of the FR band, including photons generated by transpectral processes (e.g., *(6)*), appears to be negligible at depth (Fig 3A and B). Projection of the %PrPr of DPH from different centric diatom species or environmental sequences based on their *in vitro* photochemical properties (Fig 1D, S2 and Table S2) indicated that they respond to the same environmental cues as those described above for *P. tricornutum* DPH (Fig.S7). In addition to the spectral composition, we also took light intensity into account by calculating DPH photoconversion rate constant in different environmental contexts. This revealed that DPH is sensitive enough to detect light in low light environments, as it reached photoequilibrium after 10 min as deep as 110m in clear waters (Fig.3C).

**Fig. 3.**
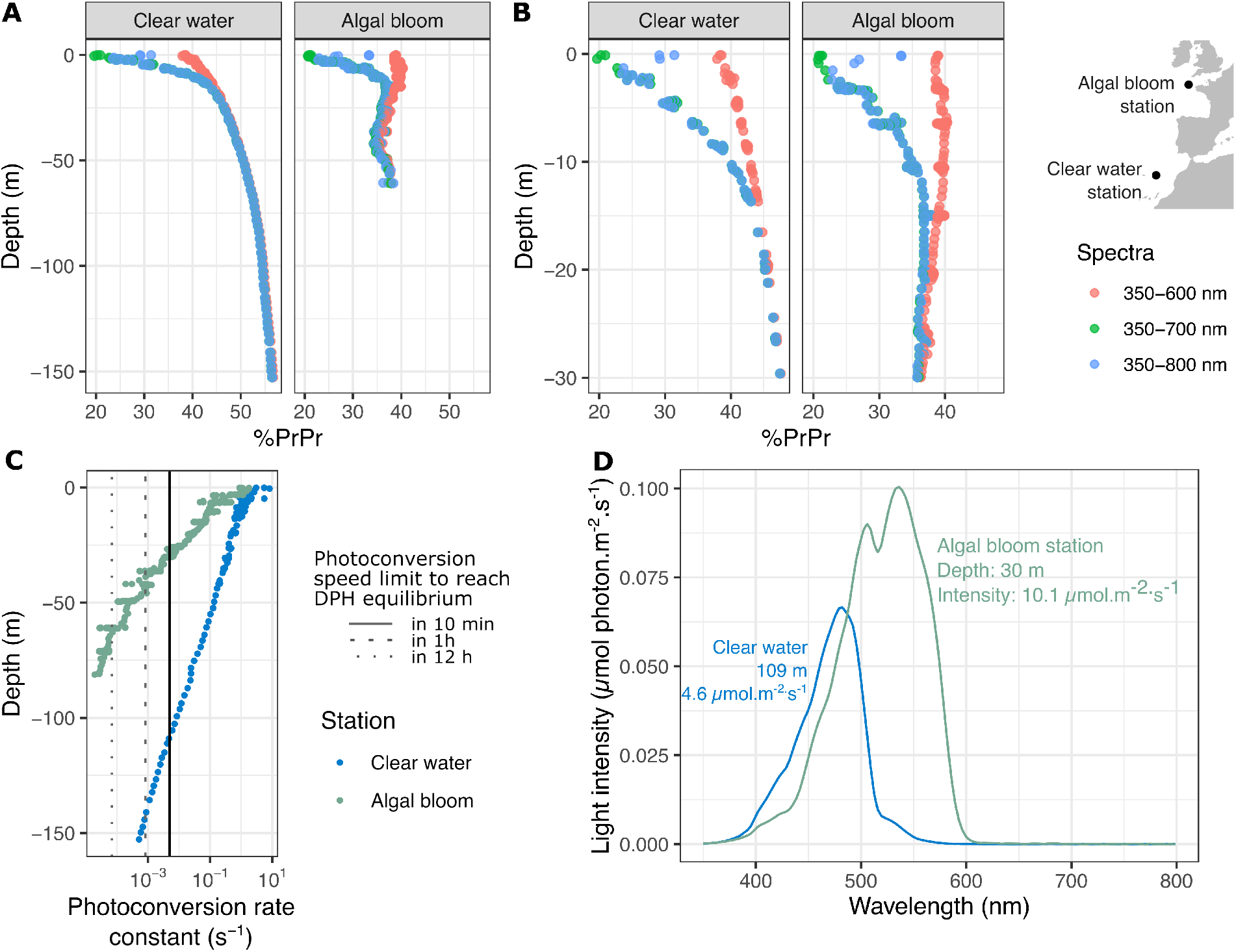
DPH integrates blue to red wavelengths and acts as a sensitive detector of depth and optical context in marine environments. (**A** and **B**) Projection of PtDPH photoequilibrium (%PrPr) in *in situ* measured marine light fields representative of two marine situations (PS71-279 and PS75-299 in *(21)*) over the whole sampled water column (A) or in the first 30 m (B). The effect of long wavebands on DPH activity was assessed by removing FR (700-800 nm) or FR and R (600-800 nm) wavebands. (**C**) PtDPH photoconversion rate constant in *in situ* light fields. Vertical lines indicate photoconversion rate constants required to reach DPH equilibrium in 12h, 1 h or 10 min. (**D**) Light field spectra at the maximal depth for which DPH is at equilibrium after 10 min.

While the function of DPH in the environment is still elusive, the remarkable response patterns shown here indicate that it may participate in the regulation of physiological plasticity in diatoms, such as photoacclimation. By adding to the information provided by light intensity (which is also the driver of chloroplast activity and activation of other photoreceptors), information on the bio-optical context, DPH permits detection of environmental changes. DPH can also work as a sensor of phytoplankton population density, drawing an interesting parallel with phytochrome-mediated neighbor perception in land plants which is however based on R/FR ratio changes *(17)*. Remarkably, DPH-containing diatoms have been found mostly in temperate and polar regions, which show strong seasonal variations of phytoplankton concentration and mixed layer depths compared to the more stable tropical regions *(18–20)*. Therefore, the presence of DPH in the marine environment could be an adaptation to the pronounced annual cycles in these regions, allowing physiological acclimation to the different seasons. Finally, the DPH response described here results from the unique interplay between the underwater visible light spectrum and the full-spectrum sensitivity of DPH. Providing the first quantitative assessment of photoreceptor activity in diverse marine environments, this study provides a rationale for the presence of a R-shifted phytochrome in a R-poor environment and reveals a new paradigm of light sensing underwater.

## Acknowledgments

We thank Richard Dorrell and Francis-André Wollman for reviewing the manscript.

## Funding

Fondation Bettencourt-Schueller (Coups d’élan pour la recherche francaise-2018) (AF)

“DYNAMO” (ANR-11-LABX-0011-01) (AF)

ANR-DFG Grant “DiaRhythm” (KR1661/20-1) (AF)

ANR ClimaClock (20-CE20-0024) (AF)

Gordon and Betty Moore Foundation (GBMF4981.01) (MJ)

CNRS (MITI interdisciplinary program) (MJ)

FFEM (Fonds Français pour l’Environnement Mondial) (CB)

French Government ‘Investissements d’Avenir’ programs OCEANOMICS (ANR-11-BTBR-0008) (CB)

FRANCE GENOMIQUE (ANR-10-INBS-09-08) (CB)

MEMO LIFE (ANR-10-LABX-54) (CB)

PSL Research University (ANR-11-IDEX-0001-02) (CB)

European Research Council (ERC) under the European Union’s Horizon 2020 research and innovation program (Diatomic; grant agreement No. 835067) (CB)

ANR ‘BrownCut’ (ANR-19-CE20-0020) (CB and AF)

ANR ‘DIM’ (ANR-21-CE02-0021) (CB)

PhD grant from the French Ministry for Higher Education, Research and Innovation (MESRI) (CD)

## Author contributions

Conceptualization: MJ, JPB, AF, MR, CB, BB

Methodology: CD, MJ, JPB, JPK, JS

Investigation: CD, MJ, JPB

Visualization: CD

Funding acquisition: MJ, AF, CB

Supervision: MJ, JPB, AF

Writing – original draft: CD

Writing – review & editing: CD, MJ, JPB, AF, MR, CB, JPK, BB

## Data and materials availability

All data are available in the main text or the supplementary materials.

## Supplementary Materials

Materials and Methods

Supplementary Text

Figs. S1 to S8

Tables S1 to S4

References (*22–40*)

Data S1

