## Supplementary methods, figures and tables for "Diatom phytochromes integrate the entire visible light spectra for photosensing in marine environments"

**The PDF file includes:**

Materials and Methods
Supplementary Text
Figs. S1 to S8
Tables S1 to S4
References (*22–40*)

**Other Supplementary Materials for this manuscript include the following:**

Data S1

### Materials and Methods

Search for *DPH* and diatom aureochrome genes in environmental sequence data

Known DPH sequences were aligned with mafft v7.4 *(22)* to generate separate Centric and Pennate HMM (Hidden Markov Models) profiles for each DPH domain, covering nearly the full-length protein. HMM searches (HMMER version 3.2; http://hmmer.org/) were performed against differents set of control sequences: protein sequences of known phytochromes from streptophytes, chlorophytes, glaucophytes, cryptophytes, bacteria, cyanobacteria, fungi and stramenopiles *(6)*, the MetDB database (https://metdb.sb-roscoff.fr/metdb/), diatom genomes and transcriptomes (see Supp Data1) and a set of proteins specific for each domain (either the proteins used in the corresponding Pfam domain alignment or if these were too few in numbers, 1000 randomly selected proteins from proteins known to possess the specific domain in InterPro). All domains with a e-value<1e-5 were retrieved, clustered at 90% identity with cd-hit *(23)* and submitted at EFI *(24)* to generate a Sequence Similarity Network (SSN). The alignment score cut-off for the SSN was chosen so that domain sequences from diatom phytochromes clustered together, not linked to other diatom proteins nor to non-diatom phytochromes (as illustrated in Fig. S8; see Table S3 for the SSN alignment score values for each domain).

The same method was applied to the aureochromes with the following modifications: only one HMM profile was generated from diatom aureochrome protein sequences, covering both the bZIP and the LOV domain; SNN being insufficient to distinguish diatom aureochromes from other aureochromes, environmental sequences were placed on an aureochrome phylogenetic tree. Briefly, sequences were aligned to the aureochrome HMM model using hmmalign and the phylogenetic tree was reconstructed with FastTree (v2.1, default settings i.e. JTT+CAT, Shimodaira-Hasegawa test with 1000 resampling, *(25)*). Only sequences branching in the diatom clades (as delimited by the sister Bolidophyceae clade) were annotated as diatom aureochromes (see FigS10)

Mapped reads to the *Tara* Oceans gene catalog (MATOU-v2) *(7,8)* sequences were retrieved from each *Tara* Oceans sample. Relative abundances were expressed as the sum of all reads mapping to centric *DPH* (or centric *AUREO*) divided by the sum of all the reads mapping to centric diatom genes. The analysis was performed on average values of the four size fractions when all were available (0.8-inf, 0.8 to 5 µm or 0.8 to 3 µm; 5 to 20 µm or 3 to 20 µm; 20 to 180 µm; 180 to 2000 µm; see FigS1 in ref 8). Environmental variables measured in situ during the *Tara* Oceans campaign are available at PANGEA *(26)*. Satellite data at *Tara* Ocean sampling stations are available at *(27)* and were used to compute intra-annual variation of optical parameters (difference between the maximum and minimum value observed throughout the year).

The relative influence of environmental parameters on *DPH* abundances were assessed with Spearman's correlations or GAMs (*mgcv* package in R *(28)*). Only variables with high explanatory power in individual GAM analysis (relative DPH abundance explained by one environmental variable, pvalue of the smooth term<0.5, explained deviance>0.35) and that did not correlate to each other (spearman's rho<0.7) were used for complex GAMs. The "proportion of explained deviance" attributed to each environmental parameter was obtained by performing the same GAM without this parameter; we then used the difference of explained deviance between full and reduced GAM as the "proportion of explained deviance", i.e. the weight of this parameter in the model.

Culture conditions

Wild-type *P. tricornutum* (Pt1 8.6; CCMP2561) cells, PtDPH-KO mutants *(6)*, and the reporter lines derived from them, *Thalassiosira pseudonana* CCMP1335, *Cyclotella cryptica* CCMP332, *Minidiscus spinulatus* RCC4659 and *Skeletonema costatum* RCC1617 were maintained in enriched artificial seawater at 19 °C under a 12L/12D regime using 50 µmol photon.$m^{-2}$.$s^{-1}$ white light (Philips TL-D De Luxe Pro 950). *Arcocellulus cornucervis* RCC2270 and *Shionodiscus bioculatus* RCC1991 were grown in enhanced artificial seawater at 4°C in 12hL/12hD cycles with white LED panel (Lumihome white LEDs). See also Table S1 for details of the strains.

Expression, purification and spectral analysis of DPH proteins

The Photosensory Module (PSM) of *DPH* genes were obtained either from previous studies (Pt-*DPH*, Tp-*DPH (6)*, cloned from cDNAs generated from cultures of *S. costatum*, *C. cryptica, M. spinulatus*, *S. biocatulatus*, *A. cornucervis* (briefly, cells were harvested by filtration, total RNA was extracted as in *(29)* and reverse-transcribed with Superscript III (ThermoFisher)), or synthetized by GenScript (USA) (for the 2 near full length sequences identified in the *Tara* Oceans gene catalog, *Minidiscus*-like and *A. cornucervis*-like) with codons optimized for *E. coli* expression. Sequences were amplified with the primers pairs indicated in the Table S4, and cloned into pET28a-HO vector generated by inserting the *Synechocystis* Heme oxygenase gene amplified from pKS270 vector *(30)* with the primer pair HO1xpET.HindIII.Fw and HO1xpET.NotI.rv, in HinDIII/NotI in the pET28a vector (Novagen). PSM sequences were expressed as N-terminal His6 tagged proteins in the E. coli BL21 (DE3) strain, using an auto-induction system *(31)*. Recombinant proteins were purified as in (6) and their absorption spectra of the recombinant proteins were measured immediately after purification on a Varian's Cary-50 spectrophotometer. Illumination with LEDs at 810, 630 and 405 nm were performed (approx. 1min illumination) to reach pure Pr spectra (after 810 illumination) or equilibria between the Pr and Pfr forms. These spectra were used as in Giraud et al (2010) *(32)* to calculate pure Pfr spectra and ratios of quantum yields.

Construction of Pt-DPH activity reporter lines

The 931-nt upstream region of the *HSF4.6a* gene (Phatr3_J49557) and the *YFP:FcpAt* fragment were amplified by PCR using the Phusion High-Fidelity DNA polymerase (Thermo Fisher, USA) from genomic DNA and the primer pairs : *Hsf4.6ap*_Fw0 and *Hsf4.6ap*-*YFP*_Rv, and from the pDEST-C-HA vector and the primer pairs : *YFP-Hsf4.6ap*_Fw and *FcpAT*_Rv0, respectively (Table S4). The two fragments were assembled by PCR using the *Hsf4.6ap*_Fw0 and *FcpAT*_Rv0, and the final product cloned into pGEM-T (Promega). Transgenic lines were obtained by biolistic co-transformation with the Phleomycin resistance plasmid pAF6 as described in *(33)*, in WT, WT (Tc) (Transformation control) and PtDPH-KO mutants obtained in *(6)*.

Experimental conditions for induction and reversion spectra

For induction and reversion spectra cells, reporter lines were maintained in continuous green light (LEDs, 520 nm, 22 µmol photon.$m^{-2}$.$s^{-1}$) with shaking (160 rpm) for at least two weeks before the experiments. On the day of the experiments, 2 or 4 L cultures at about 1.5x $10^6$ cells/mL were split into 20mL in plastic culture flasks (Corning). Flasks were irradiated one behind the other in front of an LED (about 25 flasks per LED) to generate a gradient of light due to self shading. For the reversion spectra flasks were exposed in an incubator to 10 min of far-red light (800 nm, 60 µmol photon.$m^{-2}$.$s^{-1}$) prior to illumination. The light was turned on for 10 min with no other light source then turned off, and the flasks remained in the dark for an additional 5 h 50 min time needed to have a stable amount of YFP/ cells (FigS3). Shaking was maintained during the illumination and the dark period. The same experiment was repeated for all the tested lights (LEDs center at 405, 430, 470, 730, 740, 765, 810, 850 nm for the induction fluence responses, and 405, 520, 630, 690 and 730 nm for the reversion fluence responses). Multi-wavelength expositions were performed in incubators equipped with Blue (445 nm), Green (520 nm), Red (680 nm) and FR (800 nm) LEDs. Light mixes were chosen based on the induction and reversion spectra results (ratio of wavebands and intensities) to cover ratios of DPH absorption not covered before, and to use light intensities saturating DPH reaction in 10 min. YFP measurements were performed as for the induction spectra, on cultures grown in continuous green light, splitted in 20mL flasks, exposed to 10 min of mixed illumination then transferred in the dark for 5h50 min. These experiments were done twice for each line, with two technical replicates each time.

At the end of the dark period, each flask was sampled and passed through a MAQSQant flow cytometer. The YFP signal per cell was recorded in the B1 channel (excitation 450 nm, 500 to 550 nm emission filters). 29700 events were measured, and the YFP signal per cell was averaged on the whole diatom population. The YFP signal was normalized between the signal from cells that went directly from green (growth condition) light to darkness (normalized to 0) and to cells that were exposed to 10 min of FR (765 nm) and then darkness (normalized to 1). This allowed us to compare different lines with different basal expressions of the YFP.

Light intensities and spectra were measured with a BLUE-Wave Miniature Spectrometer (StellarNet).

The different equations used for the modeling of DPH activity are issued from (13), and are presented in detail in Supplementary Text. Briefly, analysis of the normalized YFP signal at "saturating" light intensities was done for each line using the 4 flasks exposed to the highest intensities for monochromatic illumination, which was fitted with the nls function in R (Equation 6 in Supplementary Text) to estimate the ratio of quantum yield. For the analysis of the YFP-light intensity curves, for each line and for each light, the YFP signal was fitted to Equation 7, which is an exponential, with the DPH cross-section (exponential constant) as parameter to estimate.

<u>RNA extraction and gene expression analysis</u>

WT and PtDPH KO mutant lines were grown as described for the induction spectra experiments, and exposed for 30 min of saturating blue (445 nm, 30 µmol photon.$m^{-2}.s^{-1}$) or far-red (800 nm, 200 µmol photon.$m^{-2}.s^{-1}$) light, or placed in the dark. About 1.5x $10^8$ cells were harvested by filtration and total RNA was extracted as in *(29)*. For qRT-PCR, 600 ng of total RNA were reverse-transcribed using the QuantiTect Reverse Transcription Kit (Qiagen, USA) and the qPCR reaction accomplished with 12.5 ng cDNA as template, following the SsoAdvanced Universal SYBR Green Supermix instructions (Bio-Rad, USA), in a CFX 96 Real-Time Detection System (Bio-Rad). Primer sequences are indicated in Table S4. Histone H4 (*H4*, Phatr3_J34971) was used as a reference gene as in *(34)* and normalization was performed to the values before specific exposure (i.e. cells grown in continuous green light at 22 µmol photon.$m^{-2}.s^{-1}$).

<u>Pt-DPH KO mutant complementation</u>

Tp-*DPH* (Thaps3_22848) coding sequence was amplified and domesticated from *T. pseudonana* cDNA by fusing with the primer pair UNS1FL.FW and UNSXFL.RV, the PCR products obtained with the primer pairs: TpPHY.D.Fw and TpPHY.Sap1.Rv, TpPHY.Sap1.Fw and TpPHY.BsaI.Rv, TpPHY.BsaI.Fw and TpPHY.Sap2.Rv, TpPHY.Sap2.Fw and TpPHY.E.Rv (Table S4).

The 1041 nt upstream of the Pt-*DPH* start codon and the 368 nt downstream the Pt-*DPH* stop codon were amplified from *P. tricornutum* gDNA with the primer pairs: PrPtPHY.A.Fw and PrPtPHY.C.Rv, and TrPtPHY.E.Fw and TrPtPHY.F.Rv, respectively. The Tp-*DPH* domesticated coding sequence, Pt-*DPH* promoter region and terminator regions were each cloned into the pL0 vector from the modular cloning system uLoop and assembled with the Venus fluorescent protein sequence in pL1 as in *(35)*. The assembled product was amplified with the primer pair UNS1FL.FW and UNSXFL.RV, and inserted into the SmaI-linearized pUC19 vector containing the Blasticidin resistance cassette of the pPTbsr *(36)* at the Eco53KI site.

Modeled and real environmental light spectra

Measured environmental light spectra were retrieved from various studies *(21)*. We used TUV v5.3 *(37)* to compute sea-levels light spectra (direct solar and diffuse) for different sun zenith angles (downwelling irradiance only). Underwater attenuation coefficients were calculated with the formula from Morel and Maritonema, 2001 *(38)*. Light spectra were then calculated down to 100m deep for different solar zenith angles and Chlorophyll a (Chl a) concentration hypothesizing that Chl a concentration would be homogeneous in the water column. CDOM and particles *(39,40)* in addition to the model of Morel and Maritonema were also considered to mimick coastal and turbid light attenuation, but with the same effect as Chl a concentration (data not shown).

### Supplementary Text

Detailed description of DPH activity model construction based on equations described in (Mancinelli, 1994) *(13)*

Phytochrome model

Phytochrome equilibrium can be theorized as the equilibrium between the Pfr→Pr transition (rate constant k1) and the Pr→Pfr reverse reaction (rate constant k2) with the following equation:

$$PfrPfr \underset{2k1}{\overset{k2}{\leftrightharpoons}} PrPfr \underset{k1}{\overset{2k2}{\leftrightharpoons}} PrPr \quad (1)$$

The rate of phytochrome photoconversion depends on phytochrome extinction coefficient, photoconversion yield and on light intensity, so k1 and k2 can be expressed as:

$k_1 = N * \sigma_{Pfr} = N * 2.3 * \varepsilon_{Pfr} * \phi_{Pfr}$ and $k_2 = N * 2.3 * \varepsilon_{Pr} * \phi_{Pr}$

where $N$ is the fluence rate, or light intensity (µmol photon.m$^{-2}$.s$^{-1}$), and $\sigma_{Pfr}$ is the photoconversion cross-section of the Pfr→Pr reaction, $\varepsilon_{Pfr}$ is the extinction coefficient of Pfr and $\phi_{Pfr}$ the quantum yield of the Pfr→Pr reaction; $\sigma_{Pr}$ is the photoconversion cross-section of the Pr→Pfr reaction, $\varepsilon_{Pr}$ is the extinction coefficient of Pr and $\phi_{Pr}$ the quantum yield of the Pr→Pfr reaction.

Considering that the total amount of phytochrome P stays constant, i.e. synthesis and degradation are negligible, then

$P = PrPr + PrPfr + PfrPfr$ (2)

Considering PrPr as the active form and total amount of phytochrome constant, we have:

$$\frac{dPrPr}{dt} = k_1 * PrPfr - 2k_2 * PrPr \ (3)$$

At equilibrium, $\frac{dPrPr}{dt} = 0$ and :

$$\frac{PrPr_{eq}}{P} = (\frac{k_1}{k_1+k_2})^2 = \frac{1}{(1+\frac{\varepsilon_{Pr}}{\varepsilon_{Pfr}}*\frac{\phi_{Pr}}{\phi_{Pfr}})^2} = \frac{1}{(1+\frac{\varepsilon_{Pr}}{\varepsilon_{Pfr}}*\eta)^2} \ (4)$$

With $\eta = \frac{\phi_{Pr}}{\phi_{Pfr}}$

Integration of Equation 3 gives the following expression for PrPr as a function of time:

$$\frac{\mathrm{PrPr}(t)}{P} = \left(\frac{k_1}{k_1+k_2} * \frac{PfrPfr(0)}{P} + \frac{k_2}{k_1+k_2} * \frac{PrPr(0)}{P} - \frac{k_1 * k_2}{(k_1+k_2)^2}\right) e^{-2*(k_1+k_2)t} + \frac{k_1}{k_1+k_2}(\frac{PrPr(0)}{P} - \frac{PfrPfr(0)}{P} + \frac{k_2-k_1}{k_1+k_2})e^{-(k_1+k_2)t} + (\frac{k_1}{k_1+k_2})^2 \ (5)$$

Use of YFP signal as indicative of DPH activity: estimation of the ratio of quantum yield

In equation 4, $\varepsilon_{Pr}/\varepsilon_{Pfr}$ can be known from the recombinant PtDPH absorption spectra. The $\varepsilon_{Pr}/\varepsilon_{Pfr}$ ratio, in both monochromatic and bichromatic illumination, as been determined with the LED spectra with:

$$\frac{\varepsilon_{Pr,LED}}{\varepsilon_{Pfr,LED}} = \frac{\sum_{\lambda=300}^{\lambda=900} A_{Pr,\lambda} * N_\lambda}{\sum_{\lambda=300}^{\lambda=900} A_{Pfr,\lambda} * N_\lambda}$$

where $A_{Pr,\lambda}$ is the absorbance of Pr at wavelength λ measured with the recombinant protein, and $N_\lambda$ the intensity of the LED measured at this wavelength.

The PrPr/P from equation (4) can be estimated from the YFP signal at saturating light at each wavelength. Our output measure is the YFP signal for a LED at wavelength λ at saturating intensities (the 4 points at highest light intensity were used), normalized between the signal from cells that went from constant green light (520 nm, growth condition) to darkness, and the signal of cells exposed 10 min to saturating far-red light (765 nm). This is expressed as following:

$$NomalizedYFP(LED\lambda, sat) = \frac{YFP(LED\lambda, sat) - YFP(520nm)}{YFP(765nm, sat) - YFP(520nm)}$$

If we consider that the YFP is linearly linked to PrPr/P, then :

$$NormalizedYFP(LED\lambda, sat) = \frac{PrPr_{eq,LED} - PrPr_{eq,520nm}}{PrPr_{eq,765nm} - PrPr_{eq,520nm}}$$

And therefore :

$$NormalizedYFP\left(LED\lambda, sat\right) = \frac{\frac{1}{(1+\frac{\varepsilon_{Pr,LED}}{\varepsilon_{Pfr,LED}}*\eta)^2} - \frac{1}{(1+\frac{\varepsilon_{Pr,520}}{\varepsilon_{Pfr,520}}*\eta)^2}}{\frac{1}{(1+\frac{\varepsilon_{Pr,765}}{\varepsilon_{Pfr,765}}*\eta)^2} - \frac{1}{(1+\frac{\varepsilon_{Pr,520}}{\varepsilon_{Pfr,520}}*\eta)^2}} \quad (6)$$

In this equation, η is the only unknown.

To determine it, the equation was fitted with the nls function in R with η= 1 as starting value, on the experimental values obtained from monochromatic illuminations.

<u>Use of YFP signal as indicative of DPH activity: sensitivity</u>

Sensitivity is here defined as the phytochrome photoconversion cross-section σ. In theory, we have the exponential constant in equation 5

$(k_1 + k_2) = N * \left(\sigma_{Pfr} + \sigma_{Pr}\right) = N * \sigma$ and

$$\sigma = 2.3 * \varepsilon_{Pr} * \phi_{Pr} + 2.3 * \varepsilon_{Pfr} * \phi_{Pfr} = A_{Pr} * \phi'_{Pr} + A_{Pfr} * \phi'_{Pfr}$$

With $\phi'_{Pr}$ and $\phi'_{Pfr}$ corresponding to the apparent photoconversion quantum yields.

The following equation (7) was fitted to the YFP levels for each LED and each line (we used the ratio of photoconversion η estimated above for PrPr in green and far-red to correct for PrPr(0)):

$$NormalizedYFP(LED\lambda, t) = \frac{PrPr(LED\lambda,t) - PrPr_{eq,520nm}}{PrPr_{eq,765nm} - PrPr_{eq,520nm}} = \Bigg[\Bigg(\frac{A_{Pfr}\phi'_{Pfr}}{A_{Pfr}\phi'_{Pfr} + A_{Pr}\phi'_{Pr}} * \frac{PfrPfr(0)}{P} +$$

$$\frac{A_{Pr}\phi'_{Pr}}{A_{Pfr}\phi'_{Pfr} + A_{Pr}\phi'_{Pr}} * \frac{PrPr(0)}{P} - \frac{A_{Pfr}\phi'_{Pfr} * A_{Pr}\phi'_{Pr}}{\left(A_{Pfr}\phi'_{Pfr} + A_{Pr}\phi'_{Pr}\right)^2}\Bigg) e^{-2*\left(A_{Pfr}\phi'_{Pfr} + A_{Pr}\phi'_{Pr}\right)*N*t} +$$

$$\frac{A_{Pfr}\phi'_{Pfr}}{A_{Pfr}\phi'_{Pfr}+A_{Pr}\phi'_{Pr}}\left(\frac{PrPr(0)}{P}-\frac{PfrPfr(0)}{P}+\frac{A_{Pr}\phi'_{Pr}-A_{Pfr}\phi'_{Pfr}}{A_{Pfr}\phi'_{Pfr}+A_{Pr}\phi'_{Pr}}\right)e^{-\left(A_{Pfr}\phi'_{Pfr}+A_{Pr}\phi'_{Pr}\right)*N*t}+$$

$$\left.\left(\frac{A_{Pfr}\phi'_{Pfr}}{A_{Pfr}\phi'_{Pfr}+A_{Pr}\phi'_{Pr}}\right)^2-\frac{1}{\left(1+\frac{A_{Pr,520}}{A_{Pfr,520}}*\eta\right)^2}\right]\Bigg/\left[\frac{1}{(1+\frac{A_{Pr,765}}{A_{Pfr,765}}*\eta)^2}-\frac{1}{(1+\frac{A_{Pr,520}}{A_{Pfr,520}}*\eta)^2}\right]$$

(7)

where %PrPr at time 0 is either the equilibrium in green (for the induction spectra) or in far-red (for the reversion spectra), *t*, the time of illumination, i.e. 10 min, N the intensity $N = \sum_{\lambda=300}^{\lambda=900} N_\lambda$

Φ'Pr and Φ'Pfr are fitted for each LED and each line. For further analysis, we used the photoconversion cross section σ calculated from Φ'Pr and Φ'Pfr fitted values.

$$\sigma = A_{Pfr}\phi'_{Pfr} + A_{Pr}\phi'_{Pr}$$

In theory, σ should be a linear combination of ($A_{Pr}$* η +$A_{Pfr}$) (dotted lined on Fig.2C).

$$\sigma = \alpha * \left(A_{Pr,LED} * \eta + A_{Pfr,LED}\right)$$

where $A_{Pr,LED} = \frac{\sum_{\lambda=300}^{\lambda=900} A_{Pr,\lambda}*N_\lambda}{\sum_{\lambda=300}^{\lambda=900} N_\lambda}$, and $\alpha$ is a constant

Fitting the linear relationship between σ and ($A_{Pr}$* η +$A_{Pfr}$) with the lm function in R gave α=0.01618±0.00279 (adjusted $R^2$=0.3901) when considering all the data points (Figure 3C), α=0.029297±0.004329 (adjusted $R^2$=0.6154) when removing the data points above 700 nm and α=0.004667±0.000386 (adjusted $R^2$=0. 0.8633) when considering only the data points above 700 nm. Given this difference, we estimated the DPH cross section σ in the natural environment as the sum of σ below and σ above 700 nm (adjusted $R^2$= 0.6267, see also below).

From the previous equations, we can calculate $\%PrPr(t) = \frac{\mathrm{PrPr}(t)}{P} * 100$ in a given environment. The light environment is described with $I_{tot} = l * \sum_{\lambda=300}^{\lambda=900} N_\lambda$ with l the bandwidth and $N_\lambda$ the intensity at wavelength λ.

We can calculate Pr and Pfr absorption in this environment with $A_{Pr} = \frac{\sum_{300}^{\lambda} A_{Pr,\lambda}*N_\lambda}{\sum_{300}^{\lambda} N_\lambda}$, from which we calculate

$$\sigma = 0.029297 * \left(A_{Pr,\lambda<700} * \eta + A_{Pfr,\lambda<700}\right) + 0.004667 * \left(A_{Pr,\lambda>700} * \eta + A_{Pfr,\lambda>700}\right)$$

(this distinction is only relevant for real environmental data with λ from 350 to 800 nm, but not for modelled light fields, λ from 350 to 700 nm)

The ratio of absorption is calculated with $\frac{\varepsilon_{Pr}}{\varepsilon_{Pfr}} = \frac{\sum_{\lambda=300}^{\lambda=900} A_{Pr,\lambda} * N_{\lambda}}{\sum_{\lambda=300}^{\lambda=900} A_{Pfr,\lambda} * N_{\lambda}}$, from which we can calculate the DPH equilibrium:

$$\%PrPr_{eq} = \frac{1}{(1 + \frac{\varepsilon_{Pr}}{\varepsilon_{Pfr}} * \eta)^2} * 100$$

($\eta$ is either the value determined *in vivo* or *in vitro*)

We also calculated PrPr(t) with PrPr(0)=0, PfrPfr(0)=1 and

$$\frac{\mathrm{PrPr}(t)}{P} = \left(\frac{k_1}{k_1 + k_2} * \frac{PfrPfr(0)}{P} + \frac{k_2}{k_1 + k_2} * \frac{PrPr(0)}{P} - \frac{k_1 * k_2}{(k_1 + k_2)^2}\right) e^{-2*\sigma*Itot*t} + \frac{k_1}{k_1 + k_2}(\frac{PrPr(0)}{P} - \frac{PfrPfr(0)}{P} + \frac{k_2 - k_1}{k_1 + k_2})e^{-\sigma*Itot*t} + (\frac{k_1}{k_1 + k_2})^2$$

From this, we calculated $\sigma * Itot$ for which PrPr(t) ≥ 0.9 $PrPr_{eq}$ as the limit for light detection by DPH, with t=10 min, 1h or 12h.

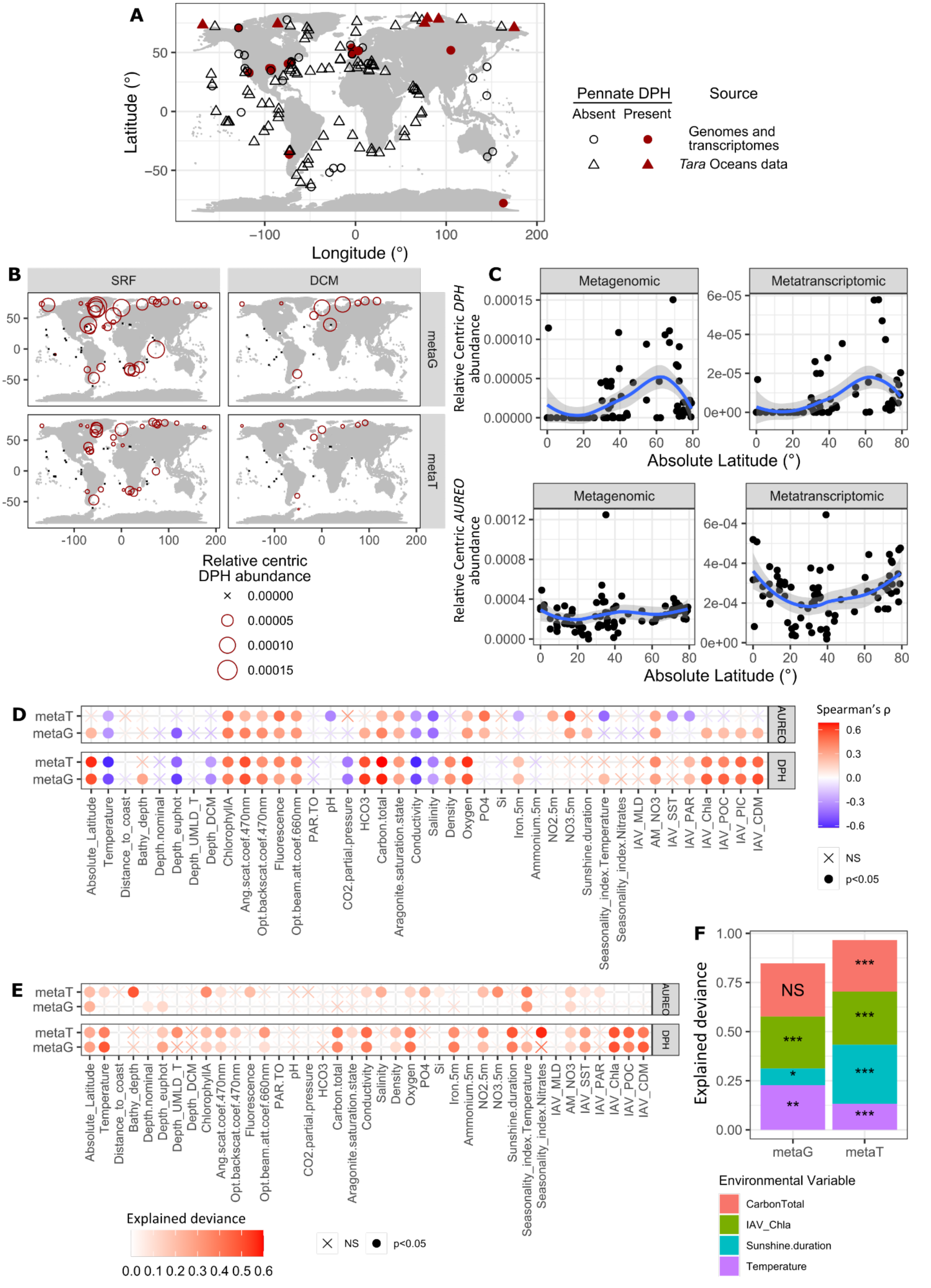
A
Latitude (°)
Longitude (°)
Pennate DPH
Absent Present
Source
Genomes and transcriptomes
*Tara* Oceans data
B
SRF
DCM
metaG
metaT
Relative centric DPH abundance
0.00000
0.00005
0.00010
0.00015
C
Metagenomic
Metatranscriptomic
Relative Centric *DPH* abundance
Relative Centric *AUREO* abundance
Absolute Latitude (°)
D
metaT
metaG
AUREO
DPH
Spearman's ρ
NS
p<0.05
Absolute_Latitude
Temperature
Distance_to_coast
Bathy_depth
Depth.nominal
Depth_euphot
Depth_UMLD_T
Depth_DCM
ChlorophyllA
Ang.scat.coef.470nm
Opt.backscat.coef.470nm
Fluorescence
Opt.beam.att.coef.660nm
PAR.TO
pH
CO2.partial.pressure
HCO3
Carbon.total
Aragonite.saturation.state
Conductivity
Salinity
Density
Oxygen
PO4
Si
Iron.5m
Ammonium.5m
NO2.5m
NO3.5m
Sunshine.duration
Seasonality_index.Temperature
Seasonality_index.Nitrates
IAV_MLD
AM_NO3
IAV_SST
IAV_PAR
IAV_Chla
IAV_POC
IAV_PIC
IAV_CDM
E
Explained deviance
F
Explained deviance
metaG
metaT
NS
Environmental Variable
CarbonTotal
IAV_Chla
Sunshine.duration
Temperature

**Fig. S1.**

*DPH* distribution is linked to latitude, temperature and optical parameters. (A) Map of the presence and absence of pennate *DPH* genes based on *Tara* Oceans sequence data and sampling location of pennate diatom strains with and without *DPH*. (B to E) Analysis of centric *DPH* biogeography from *Tara* Oceans data: (B) Map of the abundance of centric *DPH* genes and transcripts in *Tara* Oceans sampling stations (SRF: surface samples, DCM: Deep Chlorophyll Maximum; metaT: meta-transcriptomic reads, metaG: meta-genomic reads). Abundance is relative to total centric diatom genes or transcripts. (C) Centric *DPH* (top panels) and centric aureochrome (*AUREO*, bottom panels) gene or transcripts abundance relative to total centric diatom genes or transcripts as a function of absolute latitude, from metagenomic (left panels) and metatranscriptomic (right panels) data. (D) Spearman's correlation of *AUREO* or *DPH* relative abundance (in meta-transcriptomic reads, metaT, and meta-genomic reads, metaG) with different environmental parameters. (E) Generalized additive models (GAM) of *AUREO* and *DPH* relative abundances with different environmental parameters. (F) Complex GAMs explaining *DPH* relative abundance with a combination of environmental parameters.

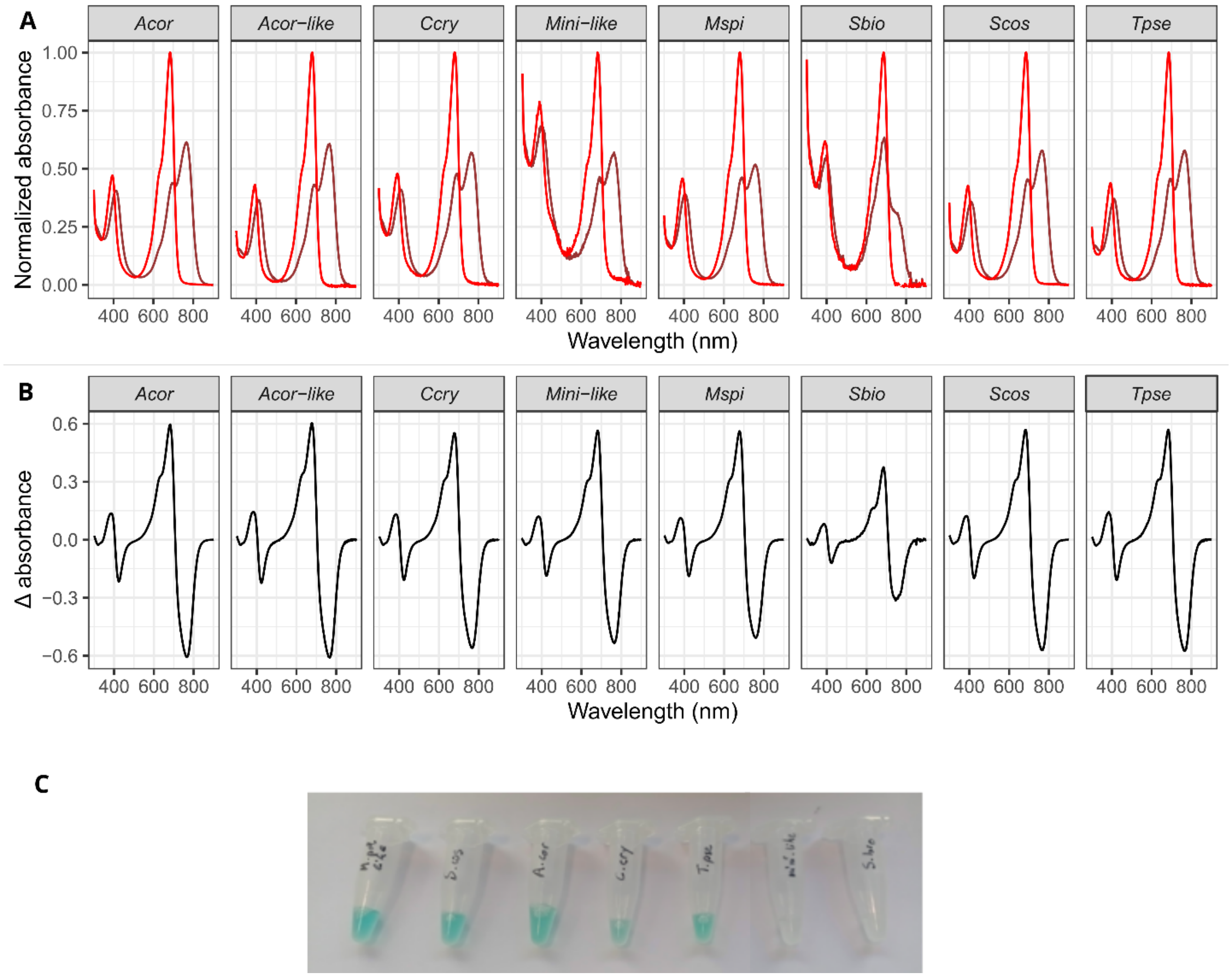


**Fig. S2.**

Spectral properties of DPH from various species are conserved. (A) Normalized absorption spectra of far-red light illuminated (red line) and red light illuminated (darker red line) recombinant photosensory domains of DPH expressed with biliverdin as the conjugate chromophore. (B) Normalized differential absorption spectra between red- and far-red- illuminated DPH. (C) Purified recombinant DPH. *Acor Arcocellulus cornucervis, Ccry Cyclotella cryptica, Mspi Minidiscus spinulatus, Sbio Shionodiscus bioculatus, Scos Skeletonema costatum, Tpse Thalassiosira pseudonana*; Mini-like: synthetic environmental sequence close to *M. spinulatus DPH*, Acor-like synthetic environmental sequence close to *A. cornucervis DPH*.

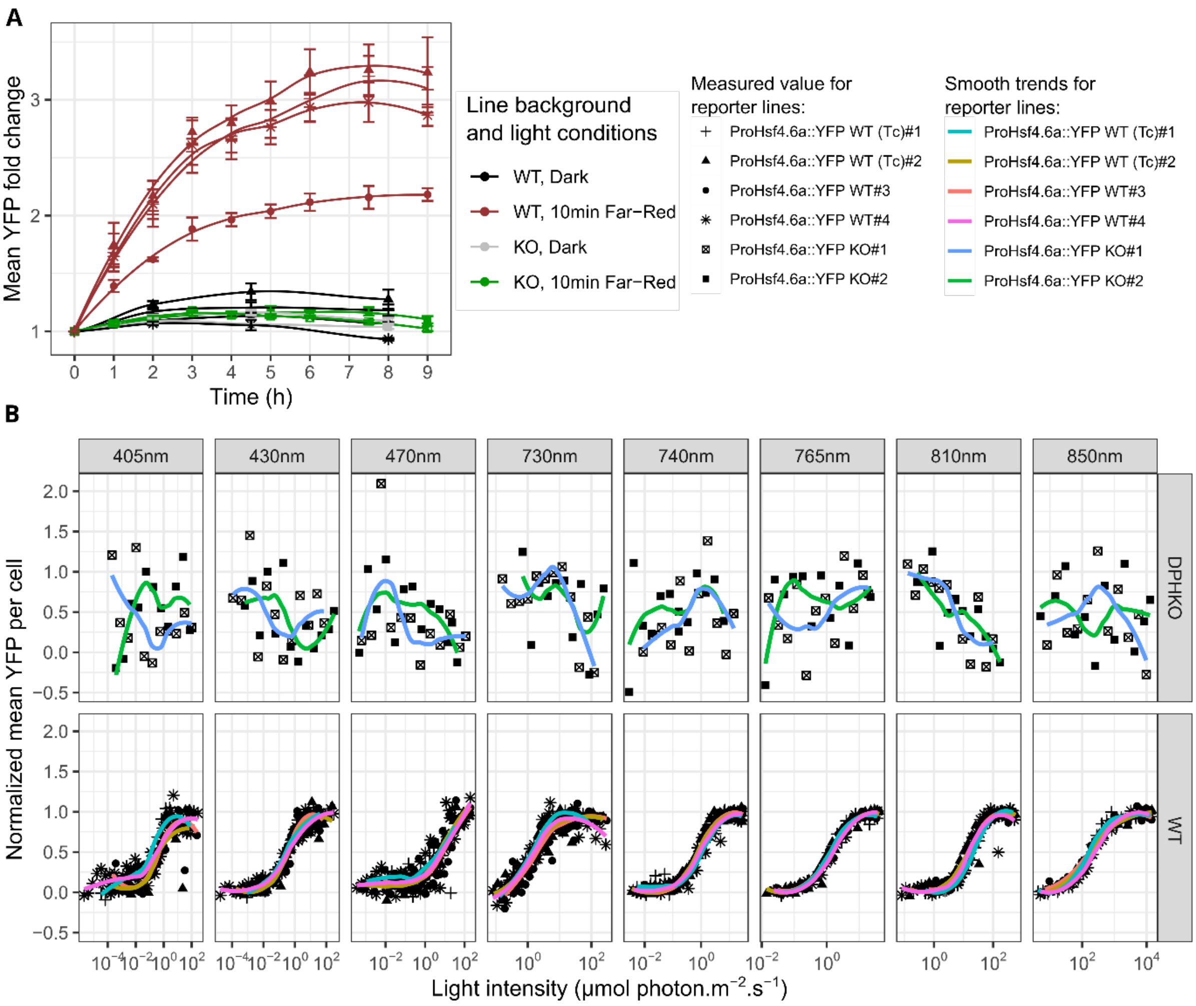


**Fig. S3.**

The PtDPH response reporter system and experimental setup. (A) Kinetics of YFP induction in different reporter line with different genetic backgrounds: *P. tricornutum* wild-type (WT), knockout PtDPH mutants (KO), or non-mutated cells originated from the same transformed colony than the KO (WT (Tc), Transformation control). Cells were grown in continuous green light (22 µmol of photon.m$^{-2}$.s$^{-1}$) and exposed for 10 min to 800 nm far-red light (800 nm, 60 µmol of photon.m$^{-2}$.s$^{-1}$) at $t_0$ then left in the dark («10 min Far-red ») up to 9 h post-irradiation, or directly transferred from green to darkness (« Dark »). YFP signals were measured by flow cytometry and normalized to $t_0$; values are mean±se of 4 replicates. (B) Fluence rate-response curves induction of YFP in response to different lights. Cells were for 10 min exposed to an intensity gradient of monochromatic lights. YFP signal was measured after 6h in the dark by flow cytometry and normalized between minimum and maximum for each light treatment and each line (mean of the 3 minimum and the 3 maximum values). WT panel are the same data as in Fig.2A with a different normalization for visualization.

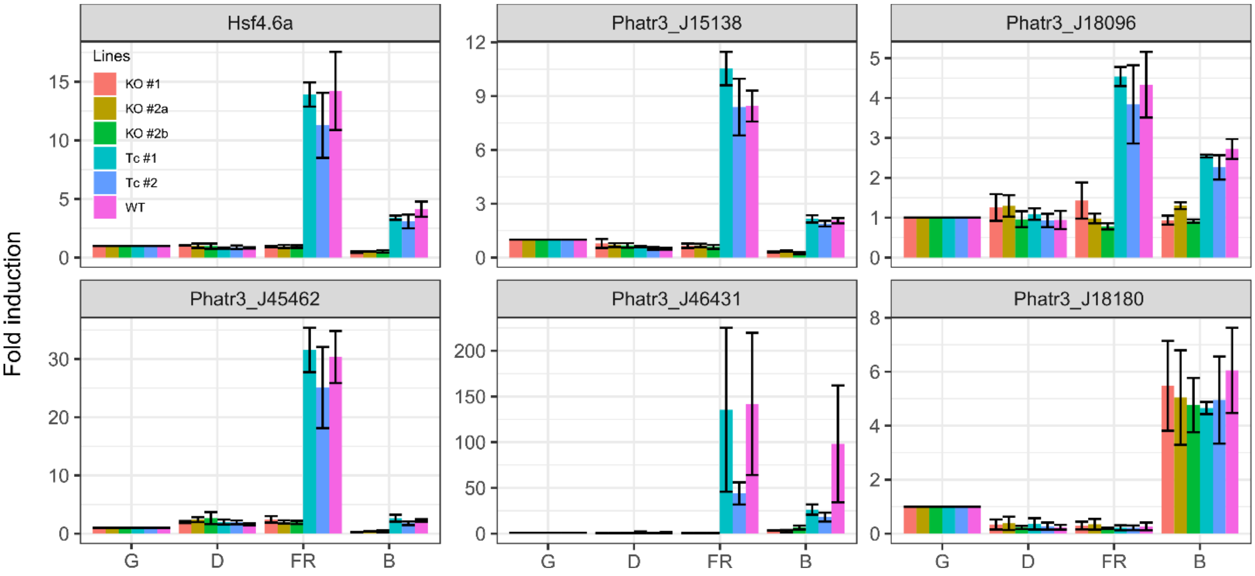


**Fig. S4.**

Expression analysis of PtDPH-regulated (*HSF4.6a*, Phatr3_J15138, Phatr3_J18096, Phatr3_J45662, Phatr3_J46431) and PtDPH-non regulated (Phatr3_J18180) genes upon far-red and blue irradiation. *P. tricornutum* WT, DPH KO mutant and their corresponding non-mutated transformant (Tc) cells were collected in their continuous green light growth condition (G) and following a 30 min of irradiation with 445 nm LED at 30 µmol photon.$m^{-2}.s^{-1}$ (B) or 800 nm LED at 200 µmol photon.$m^{-2}.s^{-1}$(FR) or kept in the dark for the same time (D). Gene expression quantifications were performed by RT-qPCR, with *H4* used as normalization and relativized to the green light condition. Values are the mean ±se on 3 independent biological replicates.

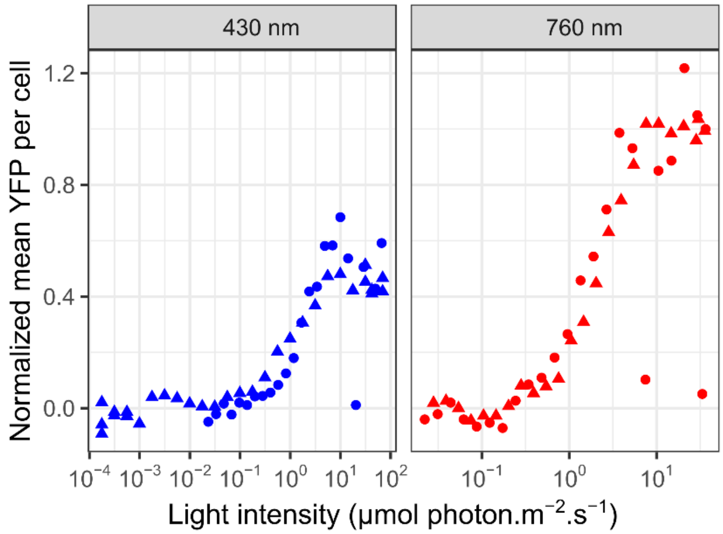


**Fig. S5.**

TpDPH restores blue and far-red-dependent YFP induction in PtDPH KO line. PtDPH KO reporter line KO #1 was transformed with Tp-*DPH* gene under the control of Pt-*DPH* promoter and terminator, and subjected to a gradient of blue (430 nm) or far-red (765 nm) lights, according to the same experimental setup used for the induction spectra. The YFP increase compared to value in the dark were normalized to the maximal YFP increase, i.e. after exposure to saturating 765 nm light. The different symbols represent data for 2 independent lines.

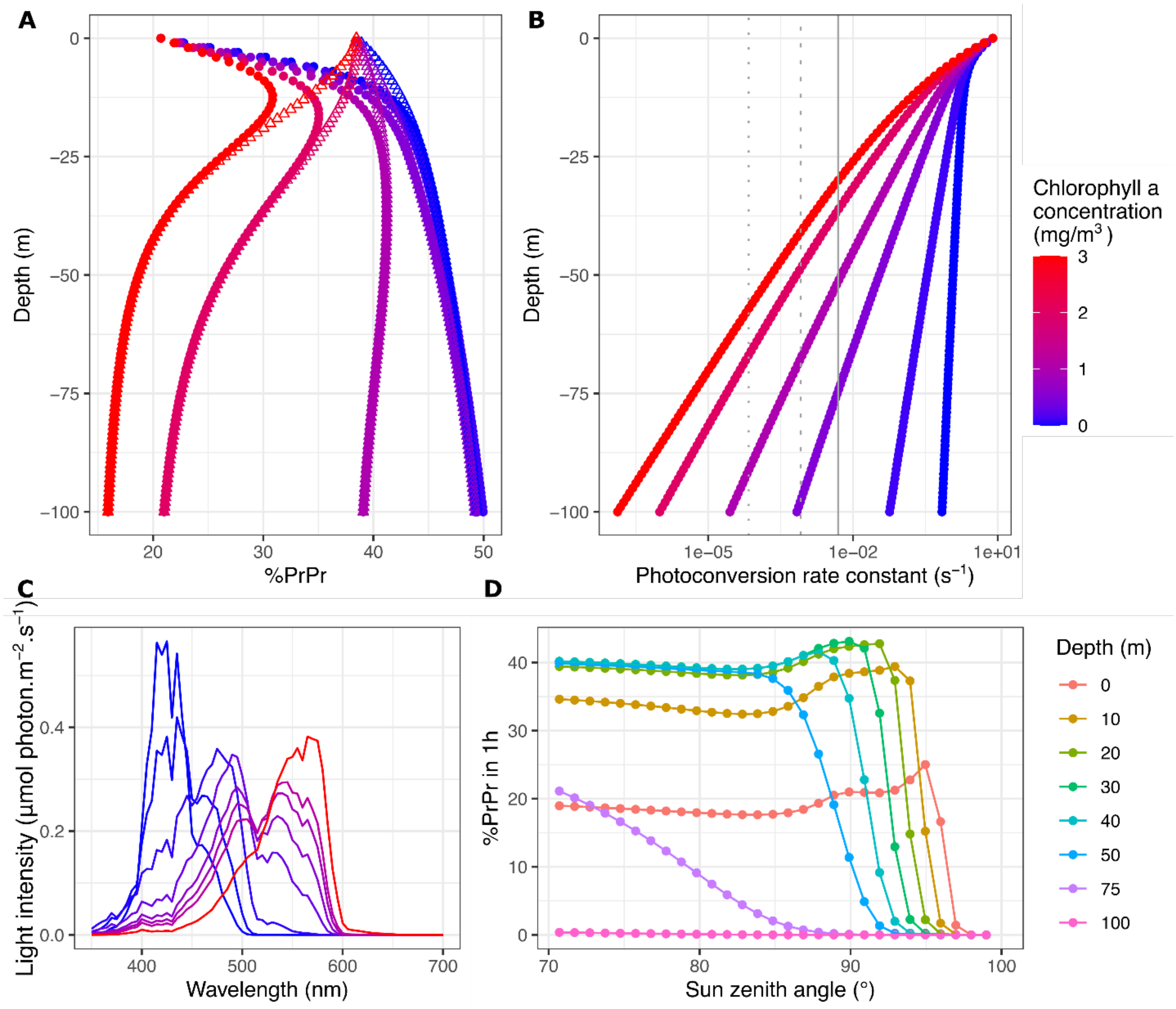


**Fig. S6.**

PtDPH activity in modeled light fields (350 to 700 nm) mimicking different underwater scenarios of phytoplankton concentration: (A) Proportion of %PrPr at depth and contribution of the red band (removing the red band, i.e. 350 to 600nm spectra, triangles), and (B) the corresponding photoconversion rate constant. Vertical lines indicate the limit for DPH to reach equilibrium in 12 h, 1 h or 10 min (from left to right). (C) Modelled light fields at the bottom of the photic zone (1% of surface irradiance) for sun at zenith. Chlorophyll a scale is the same for A, B and C. (D) DPH photoequilibrium upon 1 h of exposure of light spectrum at different depths and different solar zenith angle (Chla 1 mg/m$^3$). Note the slight increase in %PrPr around 90° at 20 to 40 m deep.

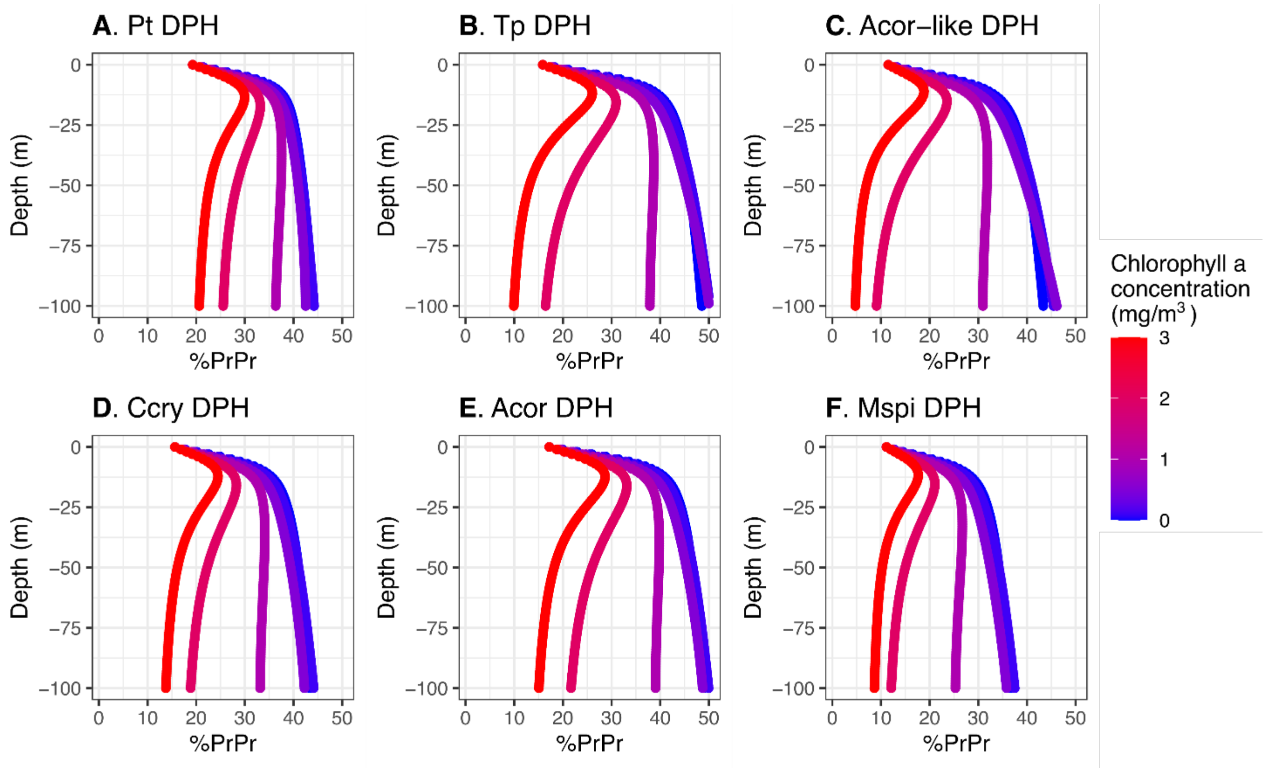


**Fig. S7.**

Projection of proportion of PrPr formed (%PrPr), in modeled light fields, with variations of chlorophyll a (Chl a) using properties of Pt DPH (A), Tp DPH (B), Acor-like synthetic DPH (C) Ccry DPH (D), Acor DPH (E) and Mspi DPH (F) determined *in vitro* from the recombinant protein absorption spectra (Table S2). The color legend is common to all panels. Note the influence of the absorption spectra (Pt DPH compared to Tp DPH) and the effect of the ratio of quantum yield (Acor DPH compared to Mspi DPH).

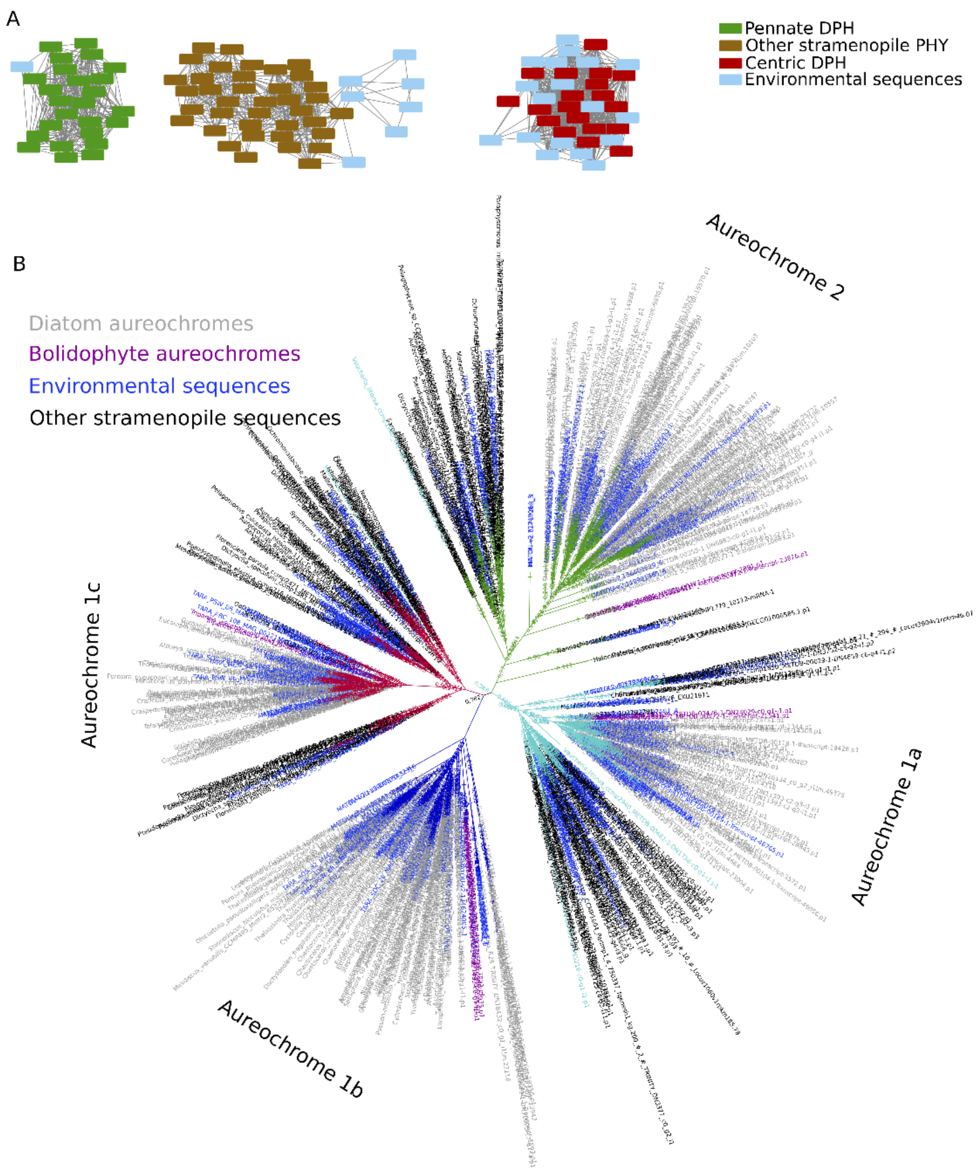


**Fig. S8.**

Method for DPH and diatom AUREO protein search. (A) Example of Sequence Similarity Network of the DPH REC domain, searched in *Tara* Oceans data showing separation of the different phytochromes (centric diatom, pennate diatom, other stramenopiles) for an alignment score threshold of 20. Environmental sequences grouping with the centric or pennate DPH will be annotated as such. (B) Phylogenetic tree of diatom AUREO search. Diatom clades are identifiable (grey label), delimited by the Bolidophyceae aureochromes (sister picoplanktonic group of diatoms, pink labels). Branches are colored by the different aureochrome types: aureochrome 1a (cyan), 1b (blue), 1c (red) and 2 (green).

**Table S1.**

Species from which DPH have been characterized in this study and previous one* from *(6)* and information about their isolation site

| Species | Strain | Sampling location, see also Data S1 |
|---|---|---|
| *Phaeodactylum tricornutum** | CCMP2561 | North Atlantic, off Blackpool, England |
| *Thalassiosira pseudonana** | CCMP1335 | North Atlantic, Moriches Bay, Forge River, Long Island, New York USA |
| *Cyclotella cryptica* | CCMP332 | North Atlantic, Martha's Vineyard, Massachusetts USA |
| *Minidiscus spinulatus* | RCC4659 | Atlantic Ocean, English Channel, Brittany coast |
| *Skeletonema costatum* | RCC1617 | Atlantic Ocean, English Channel, Brittany coast |
| *Arcocellulus cornucervis* | RCC2270 | Arctic Ocean, Beaufort Sea, at 0m depth |
| *Shionodiscus bioculatus* | RCC1991 | Arctic Ocean, Beaufort Sea, at 65m depth |

**Table S2.**

Spectral and photochemical properties of different recombinant DPH

| Phytochrome | max ΔA in B (nm) | min ΔA in B (nm) | max ΔA in R (nm) | min ΔA in FR (nm) | η |
|---|---|---|---|---|---|
| A.cor | 385 | 424 | 682 | 767 | 0.649 |
| C.cry | 384 | 424 | 677 | 768 | 0.766 |
| Minidiscus-like | 385 | 423 | 682 | 764 | NA |
| Acor-like | 384 | 424 | 678 | 768 | 0.855 |
| S.cos | 386 | 424 | 685 | 767 | NA |
| M.spi | 380 | 424 | 678 | 760 | 0.968 |
| S.bio | 386 | 424 | 685 | 746 | NA |
| T.pse* | 384 | 425 | 683 | 768 | 0.678 |
| P.tri* | 376 | 419 | 697 | 749 | 0.753 |

*absorption spectra from Fortunato et al, 2016 *(6)*. Max and min ΔA : maximum and minimum of the differential absorption spectra in the blue (B), red (R), and far-red (FR) bands. NA: not available. η were calculated with method from *(27)*.

**Table S3.**

SSN parameters for the DPH and diatom aureochrome search in *Tara* Oceans data

| Domain model | SSN alignment score threshold |
|---|---|
| Centric GAF | 30 |
| Centric PHY | 10 |
| Centric HisKA | 20 |
| Centric HATPase | 20 |
| Centric REC | 20 |
| Pennate GAF | 30 |
| Pennate PHY | 10 |
| Pennate HisKA | 20 |
| Pennate HATPase | 20 |
| Pennate REC | 20 |
| Diatom Aureochrome | 70 |

**Table S4.**

Sequences of primers used in this study

| | |
|---|---|
| qPCR-Phatr3_45662FW | CGAGGGAGCTCGGTTTATGG |
| qPCR-Phatr3_45662RV | TGATGGGAACTGTTCTGCCC |
| qPCR-Phatr3_46431FW | GGTTTGCGAGTGCATTTGGT |
| qPCR-Phatr3_45662RV | TGTCAGCAACCTCATTCCCC |
| qPCR-Phatr3_18180FW | CCGGGAACGTAGGTTTGAT |
| qPCR-Phatr3_18180RV | CCGCGGCCAACATAGCAAG |
| qPCR-H4FW | AGGTCCTTCGCGACAATATC |
| qPCR-H4RV | ACGGAATCACGAATGACGTT |
| *Hsf4.6a*p_Fw0 | TCTAGAGAGCTCGGATTTCGAATCTGTTTTGGGCA |
| *Hsf4.6a*p-*YFP*_RV | GATATCGGATCCTGTCAAAGGTTTAAGAGAATCGGC |
| *YFP-Hsf4.6a*p_Fw | AAACCTTTGACAGGATCCGATATCATGGTGAGCAAGGGCGAG |
| *FcpA*T_Rv0 | TCTAGATGAAGACGAGCTAGTGTTATTCC |
| HO1xpET.HindIII.Fw | GCAAGCTTAGAAGGAGATATACATATGAG |
| HO1xpET.NotI.Rv | GCGCGGCCGCCTAGCCTTCGGAGGTGGCGAG |
| TpPHY.D.Fw | AGGCTGTCTCGTCTCGTCTCAGGTCTCAAGGTATGAGTGTCAAAAAGAGCAC |
| TpPHY.Sap1.Rv | CAATGTGTTTCGCATGAACAGCTCTCTCGCTTG |
| TpPHY.Sap1.Fw | CAAGCGAGAGAGCTGTTCATGCGAAACACATTG |
| TpPHY.BsaI.Rv | CAAGCTGTTTATCAGGGTCACCACCCCACGTGACAC |
| TpPHY.BsaI.Fw | GTGTCACGTGGGGTGGTGACCCTGATAAACAGCTTG |
| TpPHY.Sap2.Rv | GAGATCGAACAAGTGCTCCTCAATCTGAAGGTCGTATG |
| TpPHY.Sap2.Fw | CATACGACCTTCAGATTGAGGAGCACTTGTTCGATCTC |
| TpPHY.E.Rv | TGGTAATCTATGTATCCTGTTGGTCTCTAAGCTCATCGTTCATTTTTGTGAT |
| PrPtPHY.A.Fw | GGCTGTCTCGTCTCGTCTCAGGTCTCAGGAGCCCGGGGATATCGAAGATCC |
| PrPtPHY.C.Rv | TGGTAATCTATGTATCCTGGTGGTCTCGCATTTTTAAAGGCGTGGTTCCTTG |
| TrPtPHY.E.Fw | TCGTCTCGTCTCAGGTCTCAGCTTCATGGTCGTTCATTCATAGAAG |

| | |
|---|---|
| TrPtPHY.F.Rv | TGGTAATCTATGTATCCTGGTGGTCTCAAGCGCGCTCTTTCCACCTCATCTC |
| UNS1FL.FW | CATTACTCGCATCCATTCTCAGGCTGTCTCGTCTCGTCTC |
| UNSXFL.RV | GGTGGAAGGGCTCGGAGTTGTGGTAATCTATGTATCCTGG |
| SbioPHY.NheI.Fw | GCGGCTAGCATGTCTGCCAGTTCCACCAC |
| SbioPHY_PCD.SalI.Rv | GCGGTCGACCTAAAGATTTTCCTTTTGATCTTTG |
| AcorPHY.NheI.Fw | GCGGCTAGCATGTCGGCACCTGCGGCAGC |
| AcorPHY_PCD.SacI.Rv | GCGGAGCTCCTAGTAGCTTGTTGTGTTCTCGTC |
| MspiPHY.NheI.Fw | GCGGCTAGCATGACCTCCTCCTCAACCAAC |
| MspiPHY_PCD.SacI.Rv | GCGGAGCTCCTACTTCTGATCGGCAATCAATTC |
| ScosPHY.SpeI.Fw | GCGACTAGTATGTCGTCCACCAATAGCAC |
| ScosPHY_PCD.SacI.Rv | GCGGAGCTCCTAGAGGTTTTCCTTTTGATCTTG |
| CcryPHY.NheI.Fw | GCGGCTAGCATGGCAGCACCCCACAAAAC |
| CcryPHY_PCD.SacI.Rv | GCGGAGCTCCTACTTCTGATCTTTAATCAAATC |

**Data S1. (separate file)**

Sheet 1: list of diatom genetic resources used in this study, Sheet2: Sequences of diatom phytochromes used to build hmm models, Sheet 3: Sequences of diatom aureochromes used to build hmm models. Sheet 4 and 5: Sequences of the *Tara* Oceans gene catalog (MATOU2-v2) identified as diatom aureochromes and diatom phytochromes.
